# A multiresolution framework to characterize single-cell state landscapes

**DOI:** 10.1101/746339

**Authors:** Shahin Mohammadi, Jose Davila-Velderrain, Manolis Kellis

## Abstract

Dissecting the cellular heterogeneity embedded in single-cell transcriptomic data is challenging. Although a large number of methods and approaches exist, robustly identifying underlying cell states and their associations is still a major challenge; given the nonexclusive and dynamic influence of multiple unknown sources of variability, the existence of state continuum at the time-scale of observation, and the inevitable snapshot nature of experiments. As a way to address some of these challenges, here we introduce ACTIONet, a comprehensive framework that combines archetypal analysis and network theory to provide a ready-to-use analytical approach for multiresolution single-cell state characterization. ACTIONet uses multilevel matrix decomposition and network reconstruction to simultaneously learn cell state patterns, quantify single-cell states, and reconstruct a reproducible structural representation of the transcriptional state space that is geometrically mapped to a color space. A color-enhanced quantitative view of cell states enables novel visualization, prediction, and annotation approaches. Using data from multiple tissues, organisms, and developmental conditions, we illustrate how ACTIONet facilitates the reconstruction and exploration of single-cell state landscapes.

## Introduction

Single-cell genomic technologies are revolutionizing the way tissues and cell populations are experimentally interrogated. Single-cell approaches are rapidly replacing conventional tissue-level profiling techniques, generating massive datasets in the form of cell, tissue, or organismal atlases^1–4^. Along with technological developments, single-cell biology brings new conceptual challenges. Foremost among the latter is the definition of cell identity itself, and in particular, our interpretations of a cell type and its associated dynamical states^5–7^. Technical and conceptual progress in such matters largely depends on the availability of flexible computational frameworks for efficient, meaningful, and interpretable large-scale data analysis.

In the context of single-cell transcriptomic analysis, and particularly within frameworks aiming at characterizing the structure of underlying cell states, matrix decomposition techniques are amongst the most popular approaches^8^. Although there are multiple different techniques, in essence, these methods aim to decompose a transcriptional profile into a small number of components or patterns that are presumed to optimally represent the transcriptional variability within the dataset. As part of the decomposition, the relative contribution of these patterns to the transcriptome of each cell is estimated, along with the relative contribution of genes discriminating each pattern from the others. Principal component analysis (PCA), independent component analysis (ICA), and nonnegative matrix factorization (NMF) are among the most commonly used methods; both in general applications^8,9^ and for single-cell analysis^10–14^.

Archetypal analysis (AA)^15^ is a decomposition technique that is much less frequently used, but that nonetheless offers several advantages. First and foremost, AA by design produces a more interpretable decomposition, as the underlying learned patterns are expressed as combinations of a parsimonious set of input data points^16^. In addition, AA is not constrained by certain properties of the input data, including positivity. To demonstrate the benefit of this approach for single-cell transcriptomic analysis, we recently developed the archetypal analysis for cell-type identification (ACTION) method^17^. ACTION extends AA by coupling it with separable NMF, a variant of NMF which is guaranteed to find the unique global optimum solution^18^. This coupling not only ensures the reproducibility of AA solutions, but it also improves its convergence properties and efficiency. In the context of single-cell transcriptomics, ACTION learns interpretable cell states with superior performance relative to more conventional methods^17^.

Besides matrix decomposition techniques, network-based methods are commonly used in the single-cell analysis^19–21^. In addition to a rich set of graph-based algorithms, networks bring to the realm of single-cell analysis efficient and intuitive ways to explore and visualize large-scale data. Algorithms for low-dimensional embedding^22^, community detection^23^, and diffusion-based propagation^24^ can be readily applied to visualize, group, and associate cells; provided a meaningful network-based structural representation of the cell space is available.

Aware of the complementary power of decomposition and network-based analysis frameworks, here we introduce a unique integrative way to analyze single-cell data by simultaneously learning cell state patterns, quantifying single-cell states, and reconstructing a reproducible structural representation of the underlying transcriptional state space that is geometrically mapped to a color space. By doing so, our framework allows efficient intuitive interpretation and mining of transcriptional heterogeneity at multiple consistent viewpoints: a single-cell space, a cell state space, and a gene space. Operationally, this is achieved by introducing a novel multilevel decomposition and a multiresolution cell state discovery approach that circumvents technical problems associated with transcriptomic decomposition, while accounting for a potential intrinsic biological property inherent to single-cell data: multiple meaningful levels of resolution that prohibit the specification of a single optimal number of clusters/components to partition the data. Using data from multiple tissues, organisms, and developmental conditions; we illustrate how ACTIONet can perform conventional and novel single-cell analysis tasks, including quantitative cell state and gene signature identification; cell-, state-, and gene-level visualization; gene and cell feature prediction; and automated single-cell and cell state coloring and annotation. ACTIONet and all associated downstream analysis tools are implemented in our computational environment: ACTIONet (http://compbio.mit.edu/ACTIONET).

## Results

### Overview of ACTIONet framework

The complete ACTIONet workflow consists of 5 main steps: (1) joint data transformation, dimensionality reduction, and matrix decomposition; (2) multilevel cell state representation; (3) metric space and cell similarity estimation; (4) adaptive k*nn network construction; and (5) 2/3D network embedding and de novo coloring estimation (Figure 1a).

**Figure 1.**
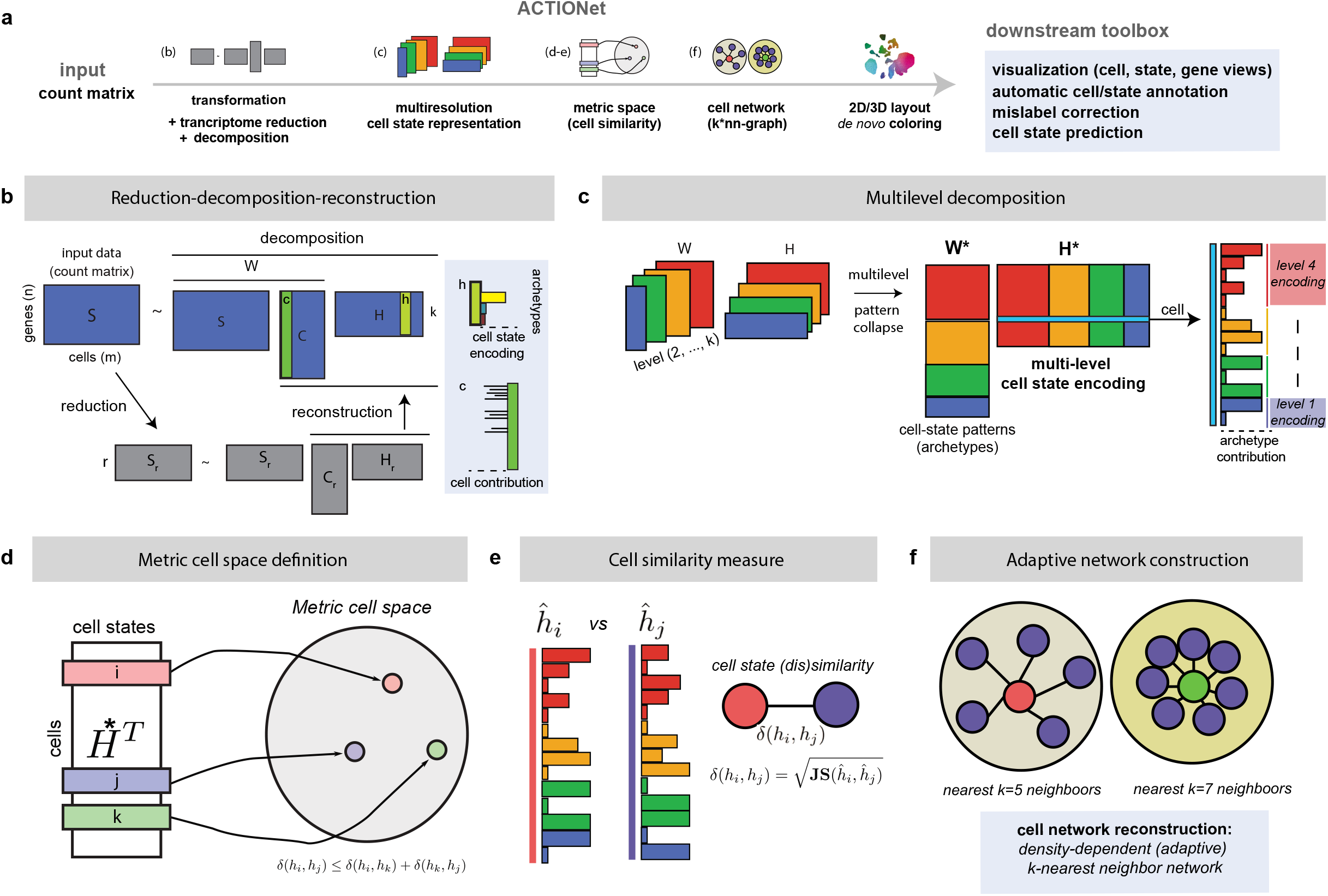
ACTIONet framework overview. **a,** Main steps in ACTIONet. **b,** ACTION-based matrix decomposition. Dimensionality reduction for feature selection (reduction) is couples with AA to perform individual-level decomposition and identify *k* latent cell state patterns (archetypes). A column *c* of the cell influence matrix C encodes the influence of cells on the patterns. A column *h* of the cell state encoding matrix H encodes the relative contribution of each pattern to the transcriptome of each cell. **c**, multilevel ACTION decomposition with an increment in the number of archetypes per level. Concatenation of individual-level matrices defines multilevel encoding (H*), cell influence (C*) and profile matrices (W*). **d-e,** Metric cell space defined by measuring distances on multilevel cell state encodings. **f,** Construction of a sparse network representation of the cell space.

Step (1) adapts archetypal analysis for interpretable matrix decomposition. Decomposition is performed over a reduced (low-rank approximation) form of the properly normalized and transformed input matrix, *f*(**S**) ~ **VS**_*r*_, where **S** ∈ genes × cells, **V** ∈ genes × *D*, and **S**_*r*_ ∈ *D* × cells represent the original expression matrix, backward linear projection matrix to the gene space, and reduced form of the transformed transcriptional profile; and *D* is the dimension of the reduced space (Figure 1b). Transformation function *f*() ensures that the inner product of columns in **S**_*r*_ approximates the original ACTION kernel, a cell similarity space^17^. Performing decomposition in a reduced space, carefully constructed to utilize the sparsity structure of **S**, ensures scalability of the algorithm (Methods). Each decomposition learns a specific number *k* of latent cell state patterns (archetypes). The relative contribution of each pattern to the transcriptome of each cell defines a cell state encoding matrix **H**, while a cell influence matrix **C** encodes the specific cells whose transcriptome jointly define each cell state pattern, here referred to as influential cells (Figure 1b). Matrices **C** and **H** are used to define, for each pattern, an expression and a signature profile matrix **W** = **SC**, and **Ŵ** = **VS**_*r*_**C**, respectively (Methods). Columns of **W** contain aggregate expression profiles of influential cells, while columns of **Ŵ** quantify the discriminating power of each gene to distinguish each pattern from the others.

The decomposition step is performed multiple times, each time increasing by one the number of cell state patterns *k* to be learned. Starting from *k* = 2, each increment determines the level of resolution at which patterns are defined. Matrix decomposition outputs are collected and reformatted in step (2), where joint, multilevel cell state encoding (**H**\*), cell influence (**C**\*), and profile (**W**\* = **SC**\*) matrices are reconstructed by concatenation (Figure 1c, Methods). Step (3) uses cell encodings in **H**\* as quantitative, low-dimensional representation of cell states to define a (multilevel) metric cell space -- i.e., space where the distance between cells can be quantitatively measured (Figure 1c,d). In step (4) the metric space is used to build a structural representation of the cell state space in the form of a graph. An efficient k*nn-algorithm^25^ automatically defines size-adaptive cell neighborhoods based on the local distribution of cell distances and density (Figure 1f). Finally, in step (5) graph layout algorithms embed the state space onto 2 and 3D spaces for visualization. To aid the latter, the 3D embedding is mapped to a perceptually uniform color space to automatically assign a color to each cell (de novo coloring). The color of the state space intuitively reflects the quantitative, gradual variability of cell states, adding information to plain 2D plots.

In addition to the multiresolution cell similarity network, the complete output of ACTIONet includes: cell state patterns; gene expression profiles and signatures for each cell state pattern; a set of dominant and nonredundant multiresolution states (see below); and a compressed cell state network. ACTIONet provides a large collection of analysis tools that make use of these outputs to facilitate the exploration and characterization of the transcriptional states in single-cell profiles. In what follows we illustrate their rationale and use.

### Level-dependent recovery of biological signals

The working principle of ACTIONet is that patterns at different resolutions are complementary, meaningful, and informative. To empirically explore the relationship between the level of resolution (*k*) and the recovery of meaningful information, we analyzed recently published data through decomposition at multiple levels of model complexity. We selected data capturing both cellular and temporal heterogeneity. We first used time-series data from mouse retinal development (n=100,831 cells)^26^, considered *k* values from 2 to 20, and analyzed the potential of the decomposition at each level to recover reported cell-type annotations (Supplementary Figure 1a) and known developmental stages (Supplementary Figure 1b). Scores of cell type and stage recovery are measured by the overrepresentation of labels in cells with high encoding on the patterns (Methods). We observed that the overall score ranking in which different cell types and developmental stages are recovered is highly variable across levels of resolution and that certain cell types/stages dominate specific but distinct decomposition levels. That is, the biological signal of different cell types and stages seems to be recovered optimally at different resolutions. We confirmed the qualitative reproducibility of these observations in multiple datasets, including Mouse Organogenesis Cell Atlas (n=100,000 cells)^27^, Mouse brain atlas (n=68,233 cells)^12,28^, Mouse Gastrulation (n=139,331 cells)^29^, Zebrafish embryogenesis, (n=38,731 cells)^30^, and human PBMC (n=7,128 cells) (10x genomics) (Supplementary Figures 1–2). In each case, we found that no single decomposition level (*k* value) seems to optimally recover the different known cell group associations. This is observed in both developmental (time-varying) datasets and in complex adult organs, and thus does not specifically reflect potential cases of a transient cell-type specification. We also confirmed that these observations are not particular results of ACTIONet decomposition. We observed qualitatively similar behavior when using ICA (Supplementary Figure 2). Our results suggest that different decomposition levels are partially informative, and therefore meaningful dominant patterns may be recovered at different levels.

### Recovered cell state patterns match cell-sorted profiles

Dominant transcriptional patterns might reflect patterns of variability other than cell type associations and time dependency. We evaluated whether cell state patterns learned through multilevel decomposition are able to recover cell-type-specific transcriptional profiles. In addition, we used independent external data to verify the interpretability of the patterns. We matched a publicly available single-cell PBMC dataset with cell-sorted, bulk RNA-seq profiles, that considers a total of 29 cell types/states (Figure 2a)^31^. To compare cell state patterns with bulk profiles, we computed partial Pearson’s correlation between ACTIONet expression profiles (**W**\*) with corresponding bulk average profiles, after adjustment for the baseline expression of genes in both profiles (Figure 2b). We found that learned dominant patterns recover transcriptional profiles consistent with expected cell-type-specific expression. We observe major correlation substructures corresponding to myeloid cells (monocytes/DCs), pDCs, B cells, T-cells, and NK cells. Among these groups, we observe the highest heterogeneity within the T-cells subgroup, which captures both its maturation state (naive/effector/memory), as well as lineage (CD4/CD8). We further verified the recovered signals using quantitative protein expression based on an available panel of antibodies against canonical markers for known PBMC subpopulations (Figure 2c). We found ACTIONet cell state pattern profiles, bulk-RNA seq, and surface marker proteins are consistent with cell states matching B-cells (CD19), Monocytes (CD14), T-cells (CD3), and NK cells (CD56). The ratio of CD45RA/CD45RO antibodies additionally verified the recovery of the maturation state for Naive and Memory T cells. Among B-cells, ACTIONet patterns successfully separate B-cells from antibody-secreting plasmablasts. pDC states resemble both myeloid and B-cells, consistent with pDC phagocytic function and lymphoid lineage association, respectively. pDC, B cell, and Mono/DC cell states cluster together, consistent with their shared antigen-presenting cell (APC) functionality. Finally, we did not find an overlap between NK cell and mature CD8 T-cell states that can be attributed to their shared cytotoxic role.

**Figure 2.**
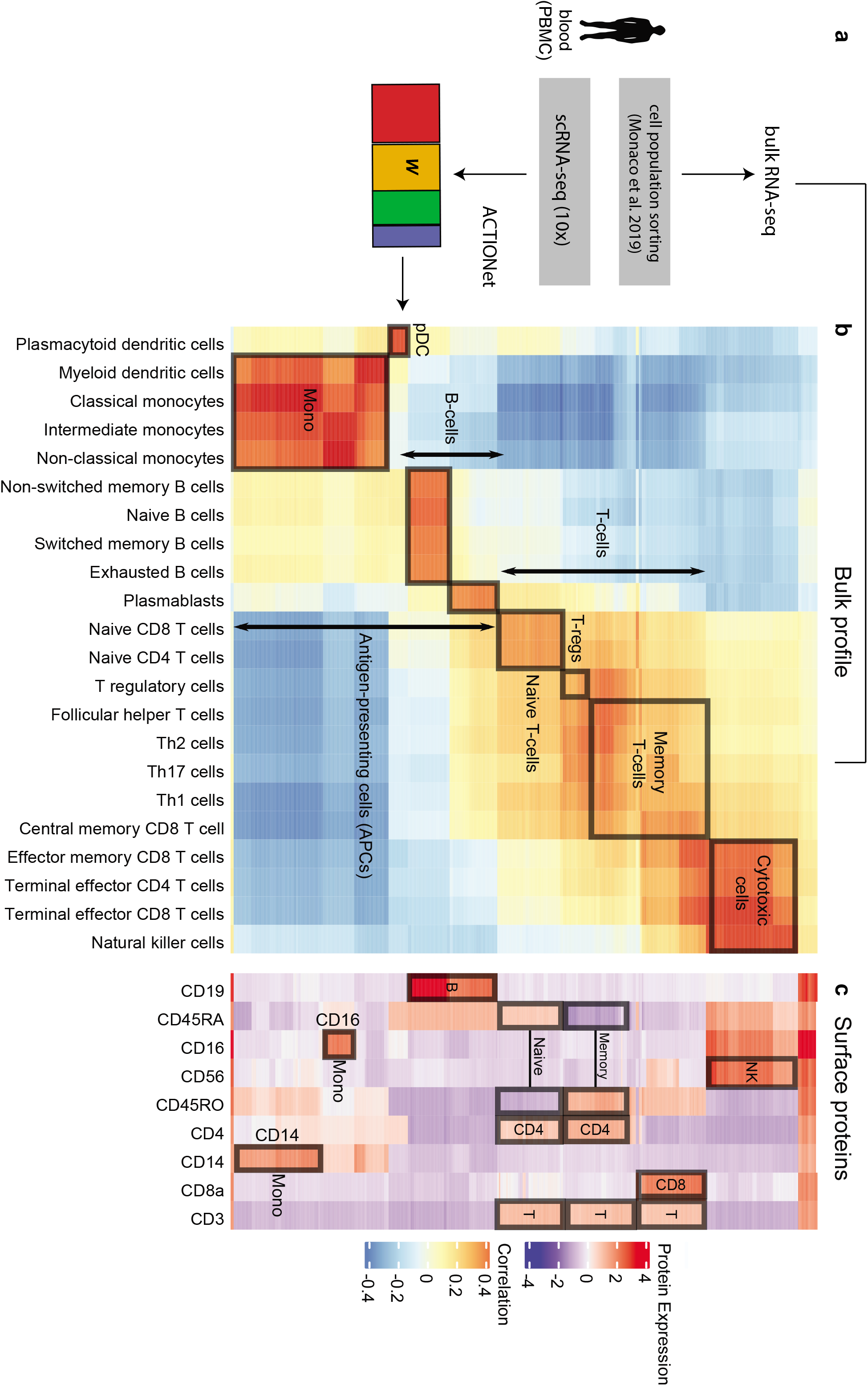
Interpretability of ACTIONet cell states. **a,** Schematic of experimental design for single-cell versus bulk profile comparison. **b**, Antibody-based measurement of characteristic marker proteins. **c**, Correlation (Pearson’s partial correlation) of cell state profiles with cell-sorted bulk RNA-seq profile.

Overall, (i) each identified pattern is highly associated with a cell type/cell state, with minimal intermixing; (ii) antibody combinations support cell annotations/association of patterns, and (iii) correlation of identified patterns with bulk, cell-sorted RNA-seq profiles allows accurate identification of both cell types and their underlying cell states.

### Structural representation of quantitative cell state space

Multilevel profiles encoded in matrix **H**\* (multilevel cell encoding) can be used as low-dimensional quantitative representations of the state of single cells. The set of encodings jointly defines an observable cell state space capturing the population’s transcriptional heterogeneity. To generate an operational cell state representation, ACTIONet defines a metric space based on multilevel cell encodings in **H**\*. To this end, columns of **H**\* are first re-scaled, to ensure each column sums to one. Given the positive nature of matrix **H**\*, each column is treated as a distribution of cells within the cell state landscape, and the square root of Jensen-Shannon divergence is used as a metric within this subspace to measure the distance between cells^32^ (Figure 1d,e).

To translate this metric space into a useful structural representation, we adapted a density-dependent nearest neighbor (k*-NN) algorithm^25^ to reconstruct a cell network. Unlike commonly used k-NN networks, k*-NN automatically identifies an optimal number of nearest neighbors for each cell, considering, in addition to proximity in our metric space, the heterogeneity and density of the neighborhood (Methods) (Figure 1f). The resulting network provides a means to visualize a large-scale state space using efficient graph layout algorithms. To directly illustrate the quantitative nature of ACTIONet cell state characterization, we introduce an automatic coloring scheme based on the projection of a 3D embedding onto a perceptually uniform color space (CIELab*). This approach effectively adds information to commonly used 2D-plots by reflecting the third dimension using colors, while linking transcriptomic with color similarity (Figure 3a). As we show below, this development also enables visually matching corresponding cell neighborhoods, underlying cell states, and gene signatures by color.

**Figure 3.**
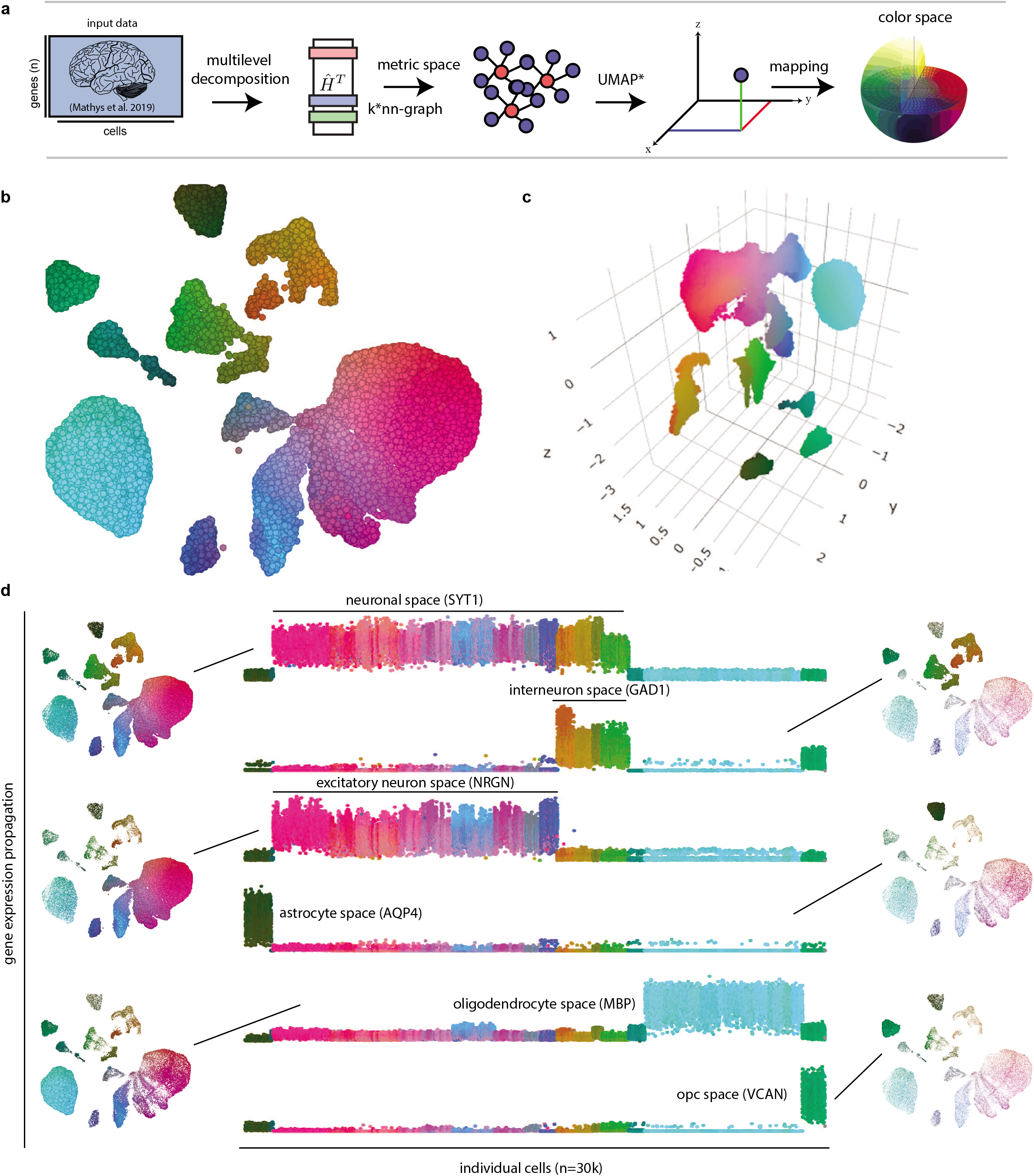
Human cortex ACTIONet cell state space. **a,** Steps used for cell state structural reconstruction and color space mapping. **b**, ACTIONet 2D visualization of the cell state and color space. **c**, 3D visualization, similar to (b). **d**, ACTIONet 1D visualization. The network propagated activity scores (vertical axes) of representative cell-type-specific markers are used to partition the cell state space, facilitating interpretation. Marker activity is mapped back to the state space (proportional to cell/node size) to aid intuition.

### Single-cell state space of the human cortex

To illustrate ACTIONet network exploratory capabilities, we first analyze the single-cell transcriptomic space of the human adult cortex. We used 35,140 cells isolated from the prefrontal cortex of 24 donors with no pathology reported previously in our recent study^33^. Along with multilevel factorization and network reconstruction, ACTIONet infers for each cell a distinctive color, which explicitly reflects its position in the 3D-embedded transcriptional state space. These colors are used to better capture cell relationships in a 2D representation (Figure 3b-c).

To explore whether cells marked by particular genes group together, ACTIONet includes diffusion-based network propagation functionality. The sparse expression values of key genes are propagated through the network, effectively inferring a smoothed expression activity profile per gene across cells (Methods). To explore how expression patterns relate to color gradients in the state space, gene activity profiles can be visualized in different ways. 1D gradient plots intuitively show how known markers partition the global state space (Figure 2d). In the human cortex space, the neuronal marker SYT1 clearly defines a neuronal space, which is further structured in inhibitory and excitatory neuronal subspaces marked respectively by exclusive GAD1 or NRGN activity. The projection of the corresponding gene activity to the 2D space (Figure 2d, right) confirms that the corresponding cells group and clearly define network neighborhoods. A similar pattern is observed for cells within the complementary glial subspace: astrocyte, oligodendrocyte, and oligodendrocyte progenitor (OPC) subspaces clearly emerge, marked respectively by AQP4, MBP, and VCAN. The 1D gradient plot representation also intuitively reflects patterns of transcriptional heterogeneity. For example, the diverse gradient of colors associated with cells showing SYT1 activity suggests that neuronal cells are the most heterogeneous, and therefore likely contain additional rich substructure. A sharp transition in color also shows that, within neurons, inhibitory and excitatory subpopulations are clearly distinct. However, relative to glial cells, the two neuronal subpopulations show increased internal color heterogeneity, a pattern likely reflecting the known abundant neuronal diversity^34^.

### ACTIONet-based automatic single cell annotation

Diffusion-based gene expression propagation can also be used to infer cell annotations. ACTIONet includes automatic cell annotation functionality that takes user-provided marker gene sets to score individual cells based on the joint activity and/or inactivity of marker genes (Figure 4a) (Methods). ACTIONet supports both positive and negative markers for cell annotation -- e.g, including genes not expected to be expressed in a given cell type. To illustrate the steps and levels of annotation enabled by ACTIONet, we use human PBMC data and consider both positive and negative markers reported recently^35^.

**Figure 4.**
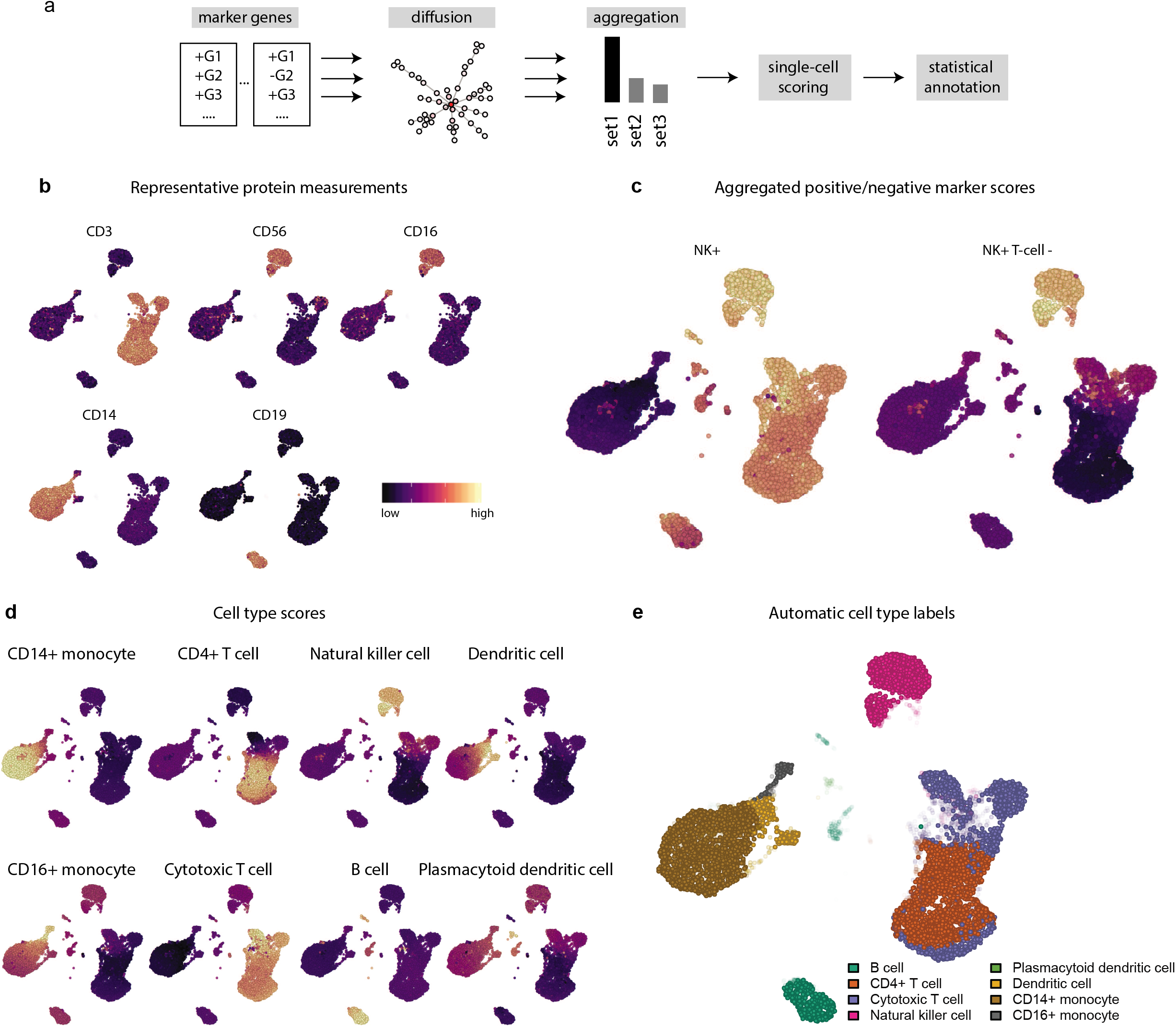
Automated cell type annotation. **a,** Steps used for automatic cell-type annotation. **b,** Projection of key surface protein expression onto the PBMC state space marks distinct cell groups. **c,** Aggregated cell-type score inferred by combining only positive (left) or positive + negative markers to automatically identify NK cell neighborhoods. **d,** Projection of aggregated inferred cell-type scores onto the PBMC state space. **e,** Automatically inferred cell-type labels for PBMC cells.

The PBMC dataset does not include reference cell annotations; however, it is accompanied by a panel of antibodies against canonical markers for known PBMC subpopulations. Figure 4b illustrates the propagation of single-cell protein antibody measurements for key markers: CD3 (T-cells), CD56 (NK cells), CD19 (B-cells), CD14 (classical monocytes), and CD16 (nonclassic monocytes/NK cells). CD16 is an example of a marker that, combined with CD56-, accurately pinpoints the CD16 monocyte population. Within the population of T-cells, CD8a distinguishes CD4 versus CD8 T-cell subtypes. These expected patterns match ACTIONet cell neighborhoods. We next use combinations of these and additional markers to automatically score and annotate the cells based on collective expression levels. We use natural killer cells to illustrate how negative markers can be used to fine-tune inferred cell-type cell scores. NK cells and memory T CD8 cells share a cytotoxic functionality, which is manifested in the aggregate score of NK-associated positive marker genes (Figure 4c, left). However, using the same NK+ markers but negatively selecting for T-cell markers (CD3D-, CD3E-’ and CD3G-), ACTIONet excludes associated T-cells, defining a purified NK cell subspace (Figure 4c, right). In addition to label assignments, the aggregate quantitative scores provided by ACTIONet can be used to analyze distribution across defined cell groups or to be projected back to the network to highlight potential cell type subspaces. Figure 4d shows that inferred cell-type scores clearly define cell subspaces for 8 major cell types. Finally, considering the profile of cell type scores computed for each cell, ACTIONet infers the most likely cell type assignment, defines a cell label, and computes an associated confidence score (Methods) (Figure 4e).

To demonstrate the generality of this approach, we independently annotated cells of the mouse^12^ and human^33^ cortex data using curated markers from two recent studies^35,36^ (Supplementary Figure 4a,b). We found strong agreement between automatic and reported annotations in both human and mouse, with mismatching observed only in cell types whose markers have been found to be transcriptionally promiscuous and/or which are prone to be experimentally mixed together (e.g., endothelial cells and pericytes) (Supplementary Figure 4c,d)^37^. When comparing group-level labels provided for mouse cells^12^, ACTIONet partitioned neurons into interneuron and excitatory subgroups and collapsed endothelial cells into a joint group (Supplementary Figure 4b).

Overall, the performance of diffusion-based cell annotation and the correspondence of annotations with well-characterized negative and positive marker gene and protein levels evidences the meaningful collective information embedded in the structure of ACTIONet.

### ACTIONet-based mislabel correction and prediction

In addition to automatic annotation based on gene expression, ACTIONet’s structure can transfer information that is only available for some cells. Using the network principle of guilt-by-association^38^, we include in ACTIONet a network-based modeling approach that predicts the most likely property of a cell, given partially known properties of its neighborhood (Methods). Using such a network-based learning framework, ACTIONet can both correct mislabels and infer missing information. We first demonstrate this functionality using cell line mixture scRNA-seq data^28^. In this data, single cells from 5 human lung cancer cell lines were experimentally mixed to provide a ground truth for benchmarking single-cell analysis. We use this gold standard dataset to test the performance of ACTIONet in correcting an incremental percentage of mislabeled cells (Figure 5a). Notably, inferred (ACTIONet corrected) labels match almost perfectly the true labels (*ARI* ~0.99) across levels of perturbation, even in the extreme case when the majority of labels (~90%) are randomly perturbed. Correct information in a fraction of cells as low as 10% percent of the total population can be successfully transferred to the whole population through collective patterns of cell network association.

**Figure 5.**
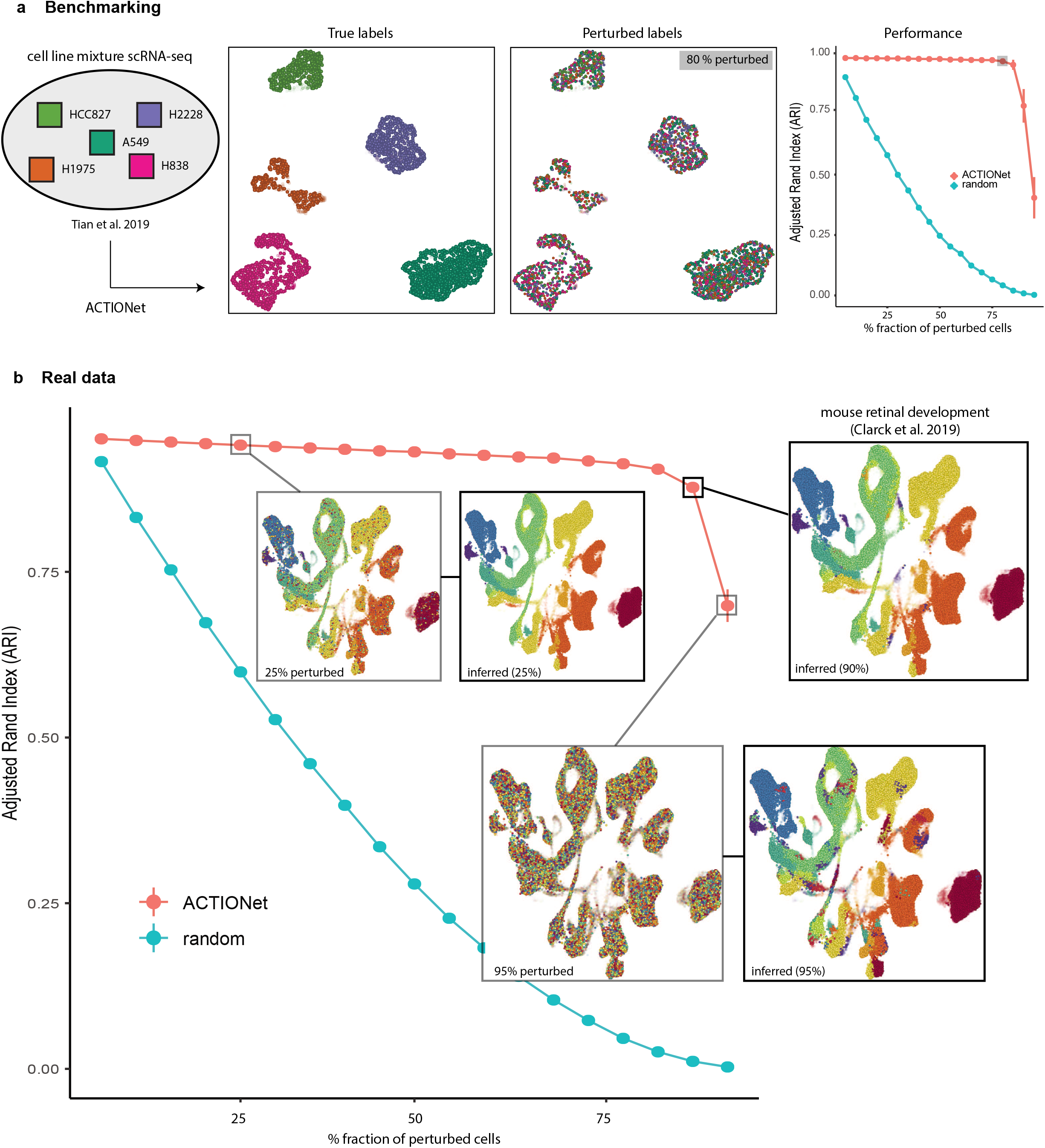
ACTIONet-based mislabel correction and prediction. **a,** Use of gold-standard cell line mixture dataset to test ACTIONet’s mislabel correction performance. An increasing fraction of the true cell labels (left) are randomly perturbed (shifted) and then corrected insing ACTIONet. Close to perfect correction performance is achieved even when perturbing 80% of the cells (right). **b,** Similar to (a), ACTIONet corrects mislabels in a real time-series dataset with more than 100k cells. The known developmental stage of the cells is randomly perturbed and then predicted.

Next, we analyzed whether a similar behavior occurs in real data, using developing mouse retina data^26^ as an example (Figure 5b). In particular, we tested whether the known developmental stage of cells can be inferred from their network context. Notably, ACTIONet performs similarly in this complex dataset, with the only substantial drop in performance occurring when 95% of the labels are randomly perturbed. Even in this extreme case, due to the coherent network structure, the information in the remaining 5% is enough to correctly infer stages for most cells (*ARI* = 0.7 ± 0.02). These results are consistent when using normalized mutual information (NMI) instead of the adjusted rand index (ARI) as a performance metric (Supplementary Figure 5).

### Identification and annotation of nonredundant multiresolution cell states

Although the full multilevel decomposition approach enhances the discovery of cell associations through network reconstruction, the independence of individual factorizations inevitably results in high redundancy: the most dominant patterns are recurrently recovered. To systematically reduce such redundancy, facilitate interpretation, and to move from a cell- to a state-level quantitative view of transcriptional heterogeneity, we utilized our cell metric space and designed an algorithm to prune multilevel patterns and identify a nonredundant, multiresolution set of cell state patterns (Methods). ACTIONet used this approach to group similar patterns across different levels into equivalent classes and, for each class, defines a representative pattern based on the lowest resolution (smallest *k*) in which a pattern of the class has been discovered. Within this framework, the size of each class provides an approximate measure of the pattern dominance. This procedure commonly reduces a multilevel space of ~200 patterns (for *k* = 2⋯20) to a multiresolution space of approximately 30 states. Annotation of these dominant patterns enables a fast and intuitive way to interpret the underlying transcriptomic states.

The dominant patterns can be mapped back to the cell state space, providing a superimposed cell state view. ACITONet’s coherent color scheme and coordinate system enable mapping cells, cell states, and gene contributions to the same space for quick, side-by-side visualization and comparison. We use PBMC data to illustrate this functionality. ACTIONet includes a cell view (Figure 6a), a cell state view (Figure 6b), and a gene view (Figure 6c). Notably, for the later, ACTIONet effectively maps informative gene signatures into a coordinate system consistent with transcriptomic heterogeneity, without directly and uniquely assigning genes to cells or states.

**Figure 6.**
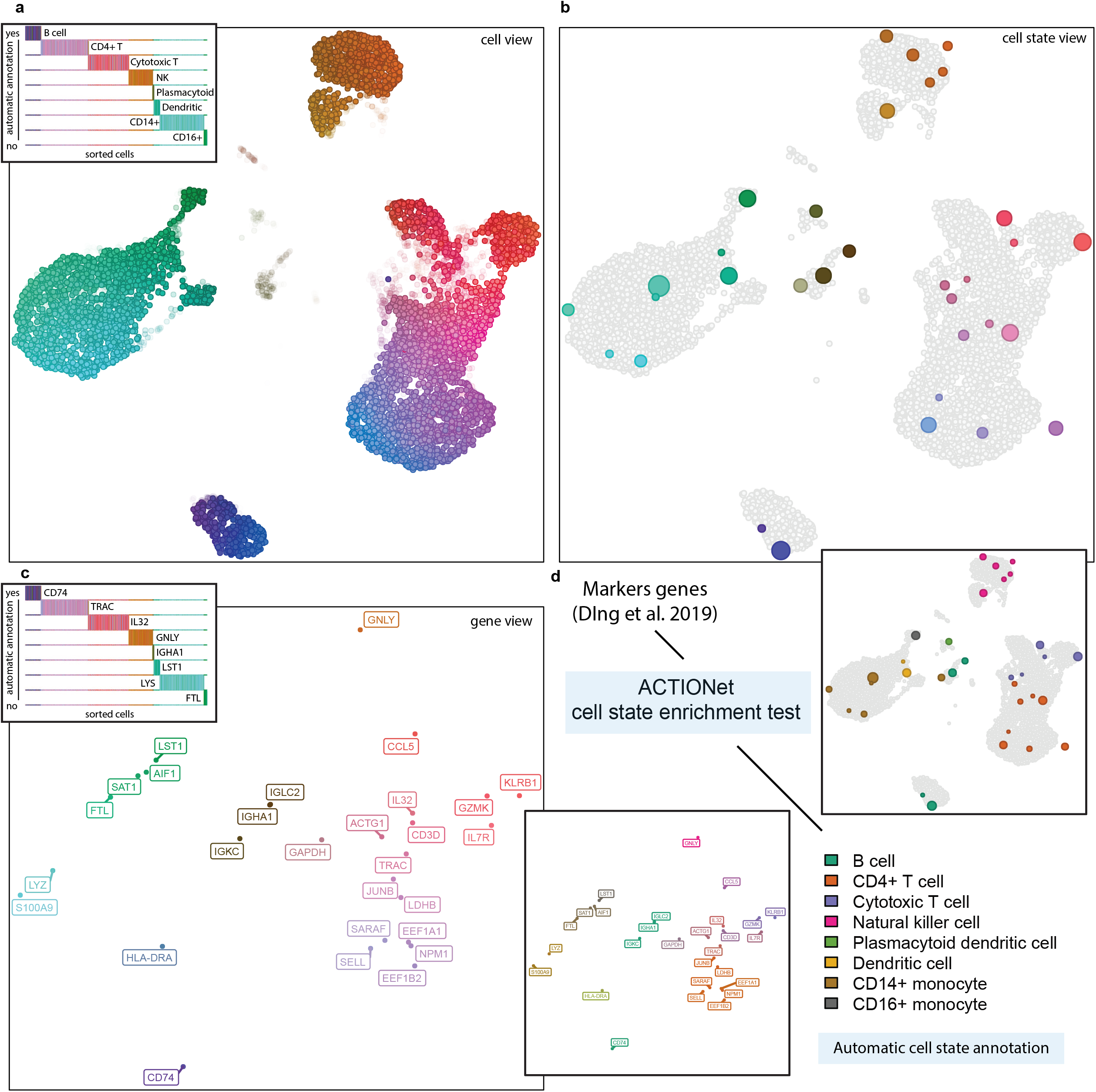
ACTIONet visualizations. **a,** ACTIONet’s cell view shows a 2D projection of all cells automatically colored by an inferred color space. A 1D plot is included to aid color interpretation based on inferred cell annotations in Figure 4. **b,** ACTIONet’s cell state view shows as colored circles the discovered dominant cell state patterns that explain transcriptional heterogeneity. Circle diameter represent the dominance of the pattern: the larger the circle the more times it was recovered at different resolutions in independent individual-level decompositions. **c,** ACTIONet’s gene view shows the signature genes that discriminate the different dominant cell states and corresponding cell neighborhoods. Matching inferred colors and coordinate system intuitively link the three views in a-c. **d,** ACTIONet’s automatic cell state annotation is used to infer for each cell state its most associated cell type using reference marker gene sets. Cell type labeling and coloring is used to aid interpretation of the different views and cell state space.

In the state space of PBMC, the cell view clearly separates neighborhoods consistent with cell types. Sorting cells based on marker-based automatic annotations, we can easily interpret the color space in terms of cell types (Figure 6a). The same coloring scheme is inherited by the dominant state patterns, which are depicted as circles in the cell state view (Figure 6b). The diameter of the circles is proportional to the dominance of the pattern; the latter quantified by the size of the equivalence class of the dominant pattern -- i.e., the number of similar multilevel patterns. The dominance is proportional to the depth of the pattern -- i.e., the level of resolution in which the dominant patterns was first detected. Each of the identified cell state patterns have a specific gene expression and gene signature profile. ACTIONet uses the latter to define the gene space and view (Methods) (Figure 6c). The color of the most influential genes again facilitates the explorative interpretation of the cell view. For a systematic interpretation, ACTIONet also includes tools to automatically annotate the cell states directly, without requiring prior annotation of individual cells.

We showed above that cell state patterns can be interpreted by direct correlation with external expression data (Figure 2). However, it is rather common to only have prior information in the form of gene sets. Certain experimental designs also enable cell labeling, for example, based on presorting experiments^31^. To further facilitate interpretation, ACTIONet includes tools to quantify the association between cell state patterns and annotations based on these two sources of information (gene sets or cell labels). To quantify gene specificity, ACTIONet uses the values of the gene signature profiles encoded in ACTIONet’s signature matrix **Ŵ**. These values effectively measure the discriminating power of each gene to discriminate a cell state pattern from the others. To quantify the preferential association of cells to cell state patterns, ACTIONet uses values of encoding matrix **H**\*. Using these quantities, efficient permutation-based enrichment analysis results in annotation matrices with enrichment scores for each cell state pattern and geneset or cell feature (Methods).

We then adopted the same set of markers previously used to automatically annotate cells (Figure 4) and this time annotate matching cell states (Figure 6d). By colors for cell type labels, ACTIONet again visually links the consistent cell-level automatic annotations (Figure 4e) and the cell state and gene views (Figure 6d).

Altogether, by combining de novo colors (Figure 7a), cell type cell-level annotations (Figure 7b), and gene set-based cell state enrichment scores (Figure 7c,d) ACTIONet can produce comprehensive annotations to aid cell state interpretation based on external prior information, including molecular pathways.

**Figure 7.**
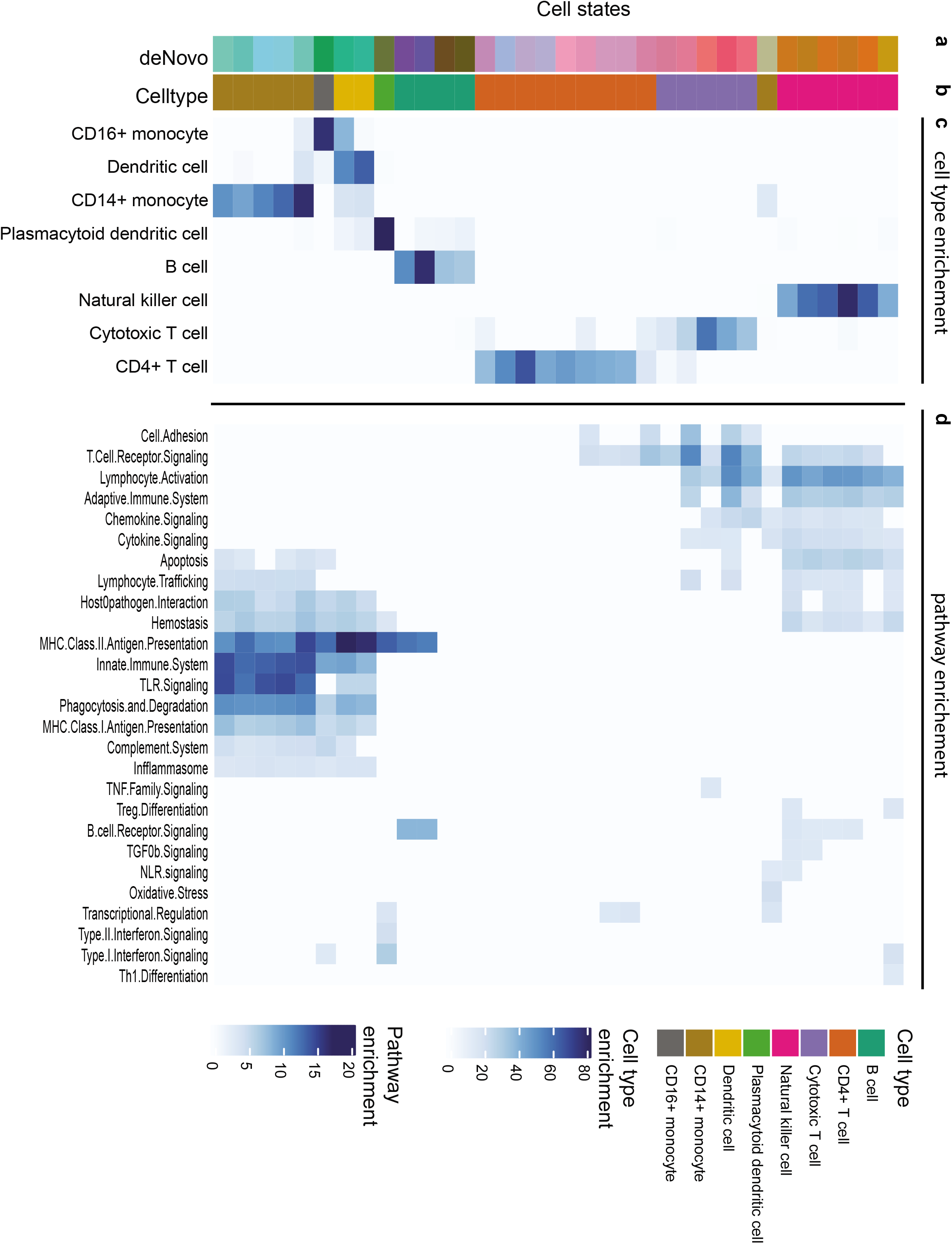
ACTIONet cell state annotation. The Heatmap shows a cell state annotation matrix. **a,** ACTIONet’s inferred de novo colors. **b,** ACTIONet’s inferred de associated cell type. **c,** Computed geneset-based cell type enrichment scores (-log p-value, permutation test). **d,** Computed geneset-based molecular pathway enrichment scores (-log p-value, permutation test). Rows correspond to dominant cell states.

### Quantitative cell state space for developmental time-series analysis

Finally, we illustrate how ACTIONet can aid the analysis and interpretation of more complex datasets, including large-scale developmental time-series. We use the state space of the mouse developing retina^26^ as an example. By combining different views of the transcriptional state space and cell state annotations, ACTIONet can provide a systematic characterization of the data. Combining the cell view, color space, and gene view, we map in one step the predominant subregions of the state space and their corresponding genetic signatures (Figure 8a,b). For example, by simple visual inspection, we automatically detect major regulators of retinal progenitor cells (RPCs) (Sfrp2, Fgf15) of neurogenic cells (Neurod4 and Neurod1), and of more neuronal Amacrine cells (Sox4, Sox11, Pax6)^26^.

**Figure 8.**
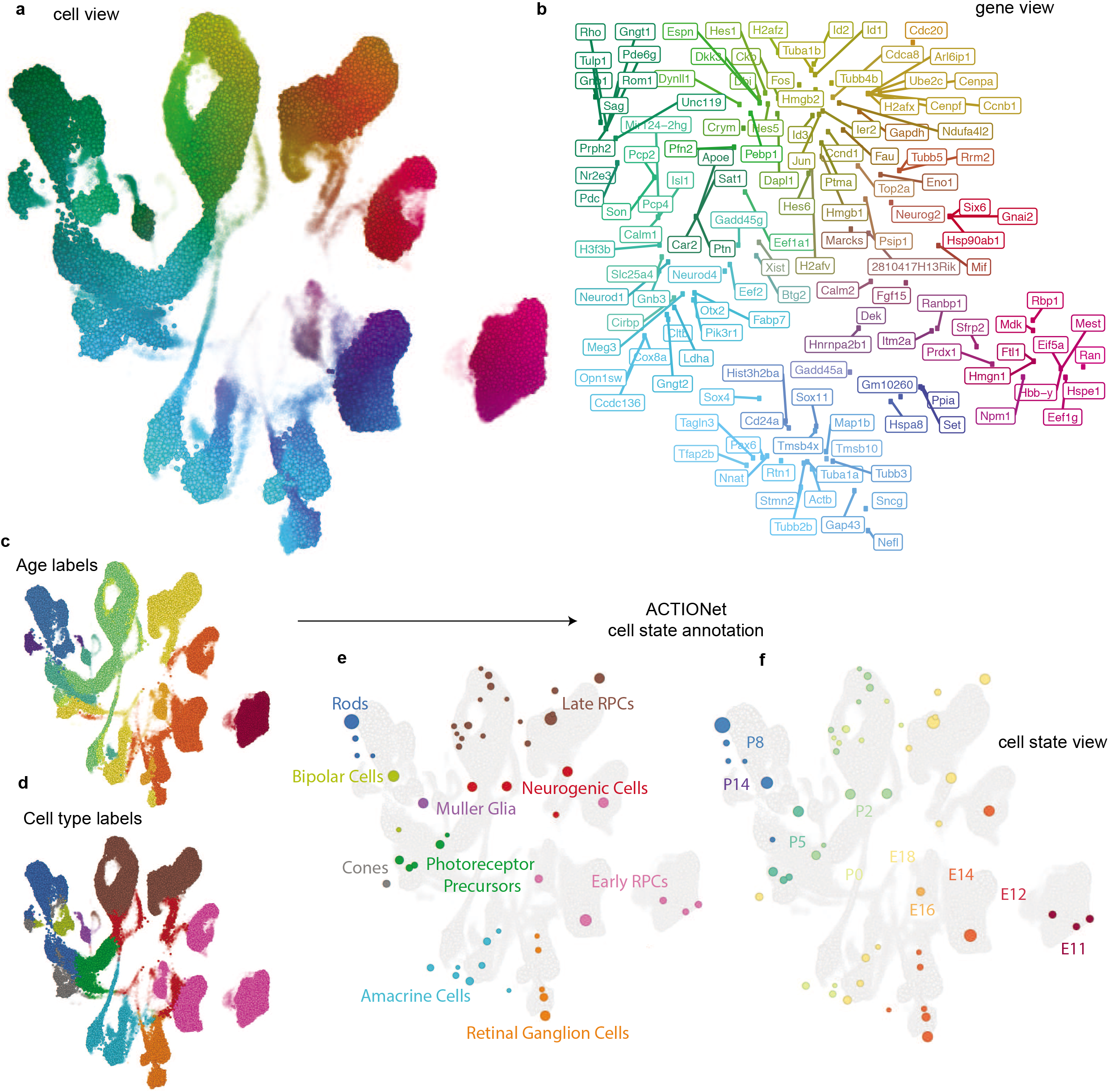
ACTIONet time-series analysis. Cell state space exploration of developing mouse retina dataset**. a,** ACTIONet’s cell view 2D state space representation. **b,** Gene-view corresponding to (a). Cell and gene views are color-coded by inferref de novo colors. **c-d,** Cell view color-coded by developmental stage (c) and cell type (d). **e-f,** Automatic cell state annotation based on cell type and developmental stage enrichment.

To visually link these exploratory observations with the available developmental stage (age) and cell type annotations, we map label-specific colors to the cell view (Figure 8c,d). This analysis clearly shows that cells aggregate in ACTIONet’s state space recovering both temporal and cell-type patterns. Finally, to more systematically analyze these patterns, we first annotate ACTIONet’s dominant cell state patterns using the cell labels and then map annotations to the cell state view (Figure 8e,f). This last analysis provides quantitative time and cell state associations to state space. Notably, the fact that ACTIONet automatically recovers a robust gradual pattern consistent with developmental time, suggests that developmental trajectories might be readily recovered through ACTIONet. And this trajectory algorithms could be directly applied to ACTIONet.

## Discussion

Using a diverse range of datasets, including multiple species and experimental designs, we show that selection of a single “best” number of archetypes in AA, or factors in general decomposition frameworks, does not optimally captures all accessible cell type, developmental, or phenotypic biological signal. Motivated by this observation, we introduced a novel framework to systematically characterize, prune, and unify the result of matrix decompositions at different levels into a nonredundant, multiresolution set of cell states that capture the complexity of the single-cell transcriptomic profiles. The end product provides a unique way to analyze single-cell states that both solves a technical problem fundamental to matrix decomposition techniques and that is connected to the intrinsic complex variability captured through single-cell profiling.

We implemented this approach, along with a large set of associated downstream analysis tools, in an easy-to-use and freely-available computational environment: ACTIONet (http://compbio.mit.edu/ACTIONet). In order to make such an approach feasible for large-scale analysis and exploration, ACTIONet includes methodological innovations at multiple steps, each of which could be independently valuable for other applications. (1) It uses a novel ACTION-based implementation of archetypal analysis for sparse matrix representation, (2) an efficient randomized SVD-based low-rank approximation (reduction step) to enable scaling with input data size, (3) an archetypal-based metric cell space construction that enables measuring cell distances, (4) an adaptive nearest-neighbor algorithm to build a multiresolution network with automatic neighbor size selection, (5) an adaptation of UMAP’s stochastic-gradient descent (SGD)-based algorithm to project a multiresolution network into 2 and 3D space, and (6) a network-based feature selection method to robustly identify a nonredundant subset of underlying cell-state patterns.

Our analyses of diverse datasets demonstrate that ACTIONet can aid the interpretation of datasets of different complexity by straightforward visualization, exploration, and statistical annotation. ACTIONet can contribute to the single-cell community both as an analysis framework to aid biological discovery, as well as a technical development to motivate novel methodologies.

## Methods

### Data sources

Cell line mixture scRNA-seq data with true labels were obtained from^28^. The dataset contains a mixture of single cells from five human lung adenocarcinoma cell lines (HCC827, H2228, H1975, H838, and A549) for a total of 3,918 cells. Human cortex data was obtained from^33^ and includes 35,140 cells profiled from 24 aged pathology-free prefrontal cortex postmortem samples. Mouse cortex data was obtained from^12^ and includes 68,233 cells profiled from the prefrontal cortex. Human peripheral blood mononuclear cell (PBMC) was obtained from the 10X Genomics website (https://support.10xgenomics.com/single-cell-gene-expression/datasets/3.0.0/pbmc_10k_protein_v3). The dataset includes 7,128 PBMC cells profiled using Total-seq protocol to simultaneously measure gene as well as cell surface protein expression (10X Genomics). Four developmental time-series datasets were downloaded from original publications. Mouse retina development data was obtained from^26^ and includes a total of 100,831 cells spanning 10 pre- and pos-natal developmental stages. Mouse organogenesis cell atlas (MOCA) was obtained from^27^. Only a sub-sample of 100,000 cells was considered. Mouse gastrulation data was obtained from^29^ and includes 139,331 cells collected from mouse embryos at nine sequential time points ranging from 6.5 to 8.5 days post-fertilization. Zebrafish developmental data were obtained from^30^ and includes 38,731 cells collected during early zebrafish embryogenesis at high temporal resolution, spanning 12 stages from the onset of zygotic transcription through early somitogenesis.

### ACTION-based transcriptome decomposition

In ACTIONet we adapted, extended, and reimplemented a scalable version of the core decomposition tool of our ACTION framework^17^. Archetypal analysis (AA) decomposition^39^ aims to identify a set of underlying transcriptional patterns (“archetypes”) whose relative contributions optimally represent the transcriptome of each cell. Archetypes, defined as a convex combination of cells, are inferred by solving the optimization problem:

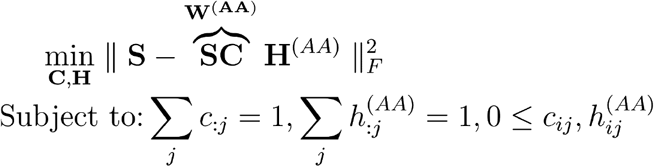

in which matrix **S** is the input profile and matrices **W** and **H** are the archetype and cell encoding matrices, respectively. While highly interpretable, AA does not guarantee convergence to the global optimum solution and its performance is dependent on the proper initialization. To remedy this issue, we first notice that separable nonnegative matrix factorization (sepNMF) can be formulated as a special case of the AA^17^:

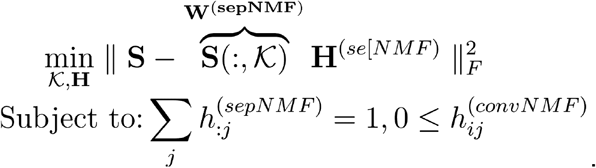

However, unlike AA, NMF can be efficiently solved using the Successive Projection Algorithm^18,40^, which has theoretical guarantees to find the global optimum solution (under a well-defined noise model). To combine the theoretical guarantees of sepNMF with the flexibility of AA, we first solve the sepNMF problem and then use the inferred factors to initialize **W**^(*AA*)^ for the AA inference problem. This process ensures the reproducibility of results and considerably enhances convergence properties. Intuitively, AA takes as input a set of pure (prototype) cells identified using sepNMF and locally adjusts these prototypes by shifting and sparsely averaging in the proximal neighborhood of the identified solutions. The enforced convexity (parsimony assumption) on the columns of **C** ensures sparsity, resulting in the definition of every archetype as a local average of a small number of cells. This approach balances the need for averaging, reduces noise, and imputes missing values while preserving subtle transitions in distinct cell states.

### Normalization and preprocessing

Prior to decomposition, ACTIONet transforms raw count data by normalizing and correcting for the baseline expression of genes. We use z-score normalization to scale each column of the input count matrix (**S**), resulting in normalized matrix **Z**. Then, we compute a baseline expression profile for each gene by averaging rows of **Z**, resulting in a vector represented by 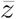. Finally, we adjust each gene for its baseline expression by orthogonalization with respect to 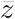,

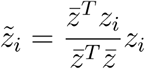

where 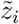 is the adjusted profile of the *i^th^* cell after correcting for the baseline expression of genes. We will represent this orthogonalization process in abstract form as 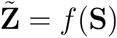. Given the rapidly growing size of single-cell datasets, computation and storage of full ACTION kernel matrix is usually infeasible. In ACTIONet, we circumvent this problem by using a reduced form of the transformed profile. To further enhance scalability, we implemented a fast randomized implementation of the SVD algorithm that has been recently developed to take advantage of the sparsity structure of the input matrix^41^. In our formulation, we use SVD to decompose the transformed, row-centered matrix, 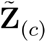 as: 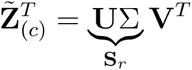.

### Multilevel decomposition and hierarchical cell similarity

Having to preselect and use only “an optimal” value of k for decomposition is a fundamental limitation of most common matrix decomposition techniques (including AA). Our observations of real single-cell data suggest that different values of *k* have different power and resolution to identify coarse vs fine-grained patterns of variability. In ACTIONet we provide a practical solution to this problem. Our solution enables gathering information across different decomposition levels. We independently perform ACTION-based decompositions, while gradually increasing the number of k archetypes. We then concatenate a posteriori all archetype profile matrices (**W**) and all cell encoding matrices (**H**) to define a multilevel cell state (archetypal) profile **W**\* and corresponding multilevel cell state encoding matrix **H**\*. We then use these two structures to reconstruct and discover multiresolution dominant patterns.

We use the multilevel profiles encoded in matrix **H**\* as low-dimensional quantitative representations of the state of single cells. The set of encodings thus jointly defines an observable cell state space. To make the theoretical construct of a cell state space of practical use, we devised a cell distance metric over cell encodings that defines an appropriate metric space. Importantly, this metric space must respect the geometric requirements of the triangle inequality. We first scale each column of **H**\* to have sum 1 and treat it as a distribution of cells in the state space of archetypes. Next, we aim to utilize normalized **H**\* to construct a metric space capturing complex relationships among cells. Let us denote with *h_i_* and *h_j_* two arbitrary columns of the normalized **H**\* matrix.

The Kullback-Leibler divergence between *h_i_* and *h_j_* is defined as:

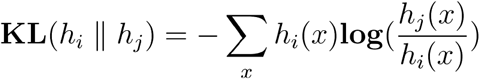

which is a positive but asymmetric measure of the difference between distributions *h_i_* and *h_j_*. We then use KL divergence to define the symmetric Jensen-Shannon divergence as follows:

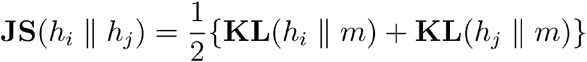

in which 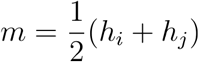. This measure is both symmetric and bounded between [0, 1], given that base 2 is used for computing the logarithm^42^. Finally, we note that JS divergence is not a metric since it does not satisfy the triangle inequality. However, it has been shown that a mono-parametric family of metrics can derive from the JS divergence. In particular, the square root of JS divergence satisfies all requirements of a metric^32^. Thus, we compute the pairwise distance between single-cell transcriptomes encodes in the corresponding columns of normalized **H**\*, as follows:

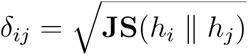

### Fast density-dependent multilevel network reconstruction

The complete cell metric space defined above provides a comprehensive operational specification of the observable cell state space. However, it is usually infeasible to store and analyze the complete distance matrix. In ACTIONet we circumvent this problem by constructing a sparse structural representation matching the metric space in the form of a graph. Given the metric nature of *δ_ij_*, we use metric (vantage point) trees to efficiently construct a nearest-neighbor graph^43^. To avoid (over)underfitting for cell types with low/high transcriptional heterogeneity, we extend conventional nearest-neighbor algorithms by adopting an adaptive strategy. We adopted an algorithm (*k*\*-nearest neighbors) originally developed for network-based regression/classification tasks^25^) that chooses the optimal number of neighbors for each node based on the distribution of neighbors’ distances. This results in a density-dependent nearest-neighbor graph. Formally, for each vertex (cell)*i*, we identify the optimal number of the neighbors to choose using the following procedure:

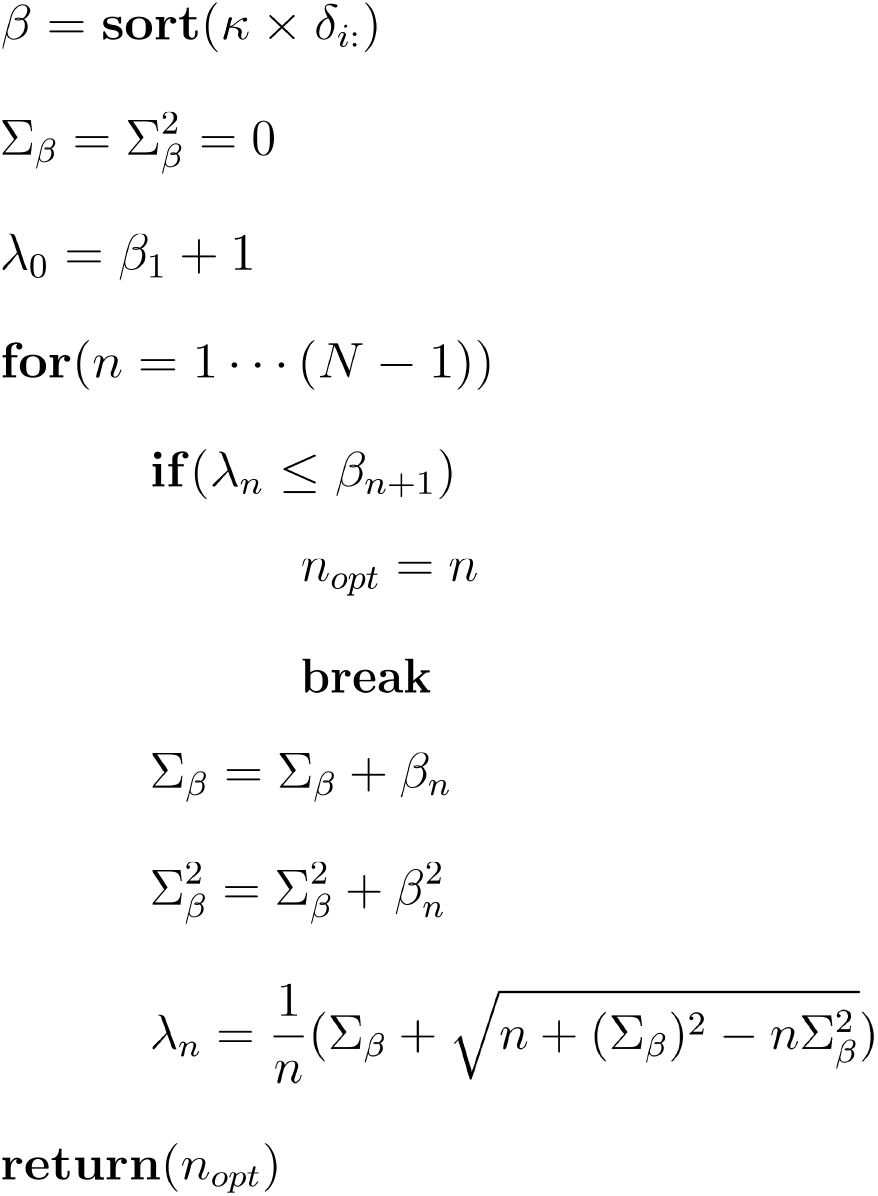

where *k* parameter adjusts the overall desired sparsity of the constructed network. Larger values of *k* result in more sparse representations, whereas smaller values of *k* result in more dense networks. In all our analyses, we use *k* = 1, that is, we use the original distances without rescaling. Finally, we note that the exact nearest-neighbor search, even in metric space, can be time-consuming for large datasets. Thus, in ACTIONet we implemented a randomized variant of nearest-neighbor search with an option to adjust the boundary of acceptable error (*ϵ*), in return for faster search^44^. When the boundary of error (*ϵ*) is set to 0, the exact search is performed.

### multiresolution cell state pattern identification

multilevel decomposition provides a comprehensive list of archetypes, each of which acts as a proxy for a potential cell state. However, two issues remain to be addressed: (i) when resolution is low (small number of total archetypes), some of the identified cell states are “too generic,” for which they represent multiple distinct cell states that are well-represented and separate at higher resolution, and (ii) same cell states can be identified/resolved in multiple levels of resolution.

To address the former issue, we first stack all *H* matrices from the multilevel decomposition, and construct the correlation matrix between rows of the stacked matrix; that is, the correlation of cell state assignments/encodings. We then construct a “backbone graph” by pruning negative correlation and using the rest of entries as edge weights linking archetypes/cell states together. In the resulting network, we then compute the weighted transitivity score^45^ for each vertex, which is a measure of connectivity within the ego-centric network of the cell state. Said differently, it indicates how often its neighbors (similar cell states) are also neighbors of each other. This will allow us to identify cell states in which are acting as “bridges” between independent cell states. To identify such cell states, we first z-score normalize the computed transitivity scores; then, we use a predefined z-score threshold to identify outlier cell states (one standard deviation below the mean is used in all experiments here).

The latter issue is more prominent in multilevel decomposition and is particularly challenging because it introduces multicollinearity in the resulting archetype matrix profile. It is also of significance as different cell types are best captured in different levels of resolution. To address this issue, we used two guiding principles: (i) each archetype/cell state is only influenced by a sparse, highly-selected number of cells. That is to say, each column of *C* matrices will have a small number of nonzero elements, that is, to our experience, independent of dataset size. We call these the “influential cells” of each archetype. (ii) for any given pair of influential cells, we can use our constructed metric space to compute their distance. Based on these two observations, we developed the following procedure to remove redundant cell states from our datasets. First, we identify the influential cells of each archetype and construct a population of all influential cells. Then, we compute all pairwise distances between these influential cells. Starting with a set of cell states, each with its own set of influential cells, we first compute the distribution of intra-cell-state cell distances, as well as inter-cell-state cell distances for every pair of cell states. We then use the t-test to compare these two distributions for each pair of cell states and iteratively merge cell states that their intra-vs-inter cell distance distributions do not significantly deviate from each other. That is to say, for a pair of cell states that the distance of cells within each cell state compared to the distance of cells between cell states are not significantly different from each other. We iteratively merge cell states, starting from the closest pairs, and their corresponding influential cells till we reach a pair of cell states that are significantly different from each other, inferred from distances of their influential cells.

### ACTIONet data visualization

ACTIONet provides a powerful computational tool amenable to well-established network analysis techniques. For efficient graph visualization, we implement low dimensional graph embedding in 2 and 3 dimensions, while preserving the underlying manifold structure. We partially adopted, slightly modified, and reimplemented the manifold learning technique Uniform Manifold Approximation and Projection (UMAP), which preserves both global and local topological features in the reduced space^22,46^. We only adopted the embedding stage of the UMAP algorithm to layout the density-dependent multilevel ACTIONet graph.

Intuitively, UMAP efficiently approximates a force-directed layout algorithm. At its core, it aims to minimize the cross-entropy of two fuzzy sets (1-simplices), one of which captures relationships between cells in the original graph and the other within the lower dimensional Euclidean subspace. The cross-entropy objective can be written as the sum of an entropy term within the projected subspace, coupled with a KL divergence term that penalizes for the deviation of point distribution from the original edge weights. These components effectively define the repulsive and attractive forces of the layout algorithm. More formally, in ACTIONet we implemented the Stochastic Gradient Descent (SGD) algorithm of UMAP to efficiently optimize the following objective function:

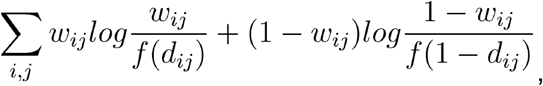

where *w_ij_* is affinity scores corresponding to the edge weights of ACTIONet graph, *d_ij_* are the euclidean distances between embedded points, and the function *f*() is defined as:

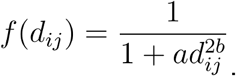

In this formulation, parameters *a* and *b* are inferred through nonlinear least-squares fitting and can be used to control the compactness of the final embedding. By testing a wide range of values, we found that these parameters behave similarly in particular regimes. To simplify the interaction with the algorithm, we precomputed a set of *a* and *b* parameters and replaced them with a new “compactness” parameter, which takes values between 0-100 that provide a practically useful range of gradually increasing compact representations for single-cell datasets.

Finally, we observed that UMAP performance improves by using “smoothed” weights computed as follows:

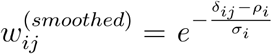

where *ρ_i_* is the distance to the closest neighbor of node *i*, and *σ_i_* is set to be the value such that:

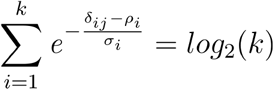

where *k* is the number of the nearest-neighbors of node *i*. We use this latter procedure in ACTIONet to convert metric distances between cells into edge weights.

### De novo coloring

We observed that a three-dimensional embedded space usually provides a very good approximation to the apparent structure of the cell state space, a feature that is sometimes misleadingly lost in the more standard 2D visualization. To directly link the two representations, while highlighting the quantitative nature of the cell state characterization enabled by ACTIONet, we introduce an automatic (de novo) coloring scheme. We first construct both 2D and 3D embeddings and use properly scaled 3D coordinates to map each cell to a color space. We adopt the CIE L*a*b* (CIELAB) color space which, unlike RGB space, is perceptually uniform: i.e, it is designed such that points with similar distances are visually perceived as having similar color differences^47^. This procedure allows us to intuitively map the three-dimensional embedding of the ACTIONet graph into a colorspace that recapitulates its overall topology. By using these colors in multiple visualizations, ACTIONet adds information about the quantitative state of cells while linking 1- and 2D cell, state, and gene views.

### ACTIONet automatic cell annotation

Our cell annotation framework uses known marker genes, both positive and negative markers, to infer the most likely cell type for each cell, individually. First, it uses a network-based diffusion method, with the ACTIONet graph as input, to impute the expression of every marker gene. Then, it computes the signed average of the imputed expression vectors to compute a score for each cell type/cell pair. Finally, it assesses the significance of these cell type-association scores by permutation test; it samples the same number of imputed genes selected at random and constructs a null model for their corresponding weighted average statistics. Then, it uses this null model to assign a z-score to each cell-type/cell association.

### Network propagation-based mislabel correction

We use a variant of label-propagation algorithm to update a given set of labels using the topology of the ACTIONet graph. In summary, we aggregate labels within the neighborhood of each cell and identify the most, neighborhood-inferred label for each cell, together with a confidence score. If the neighborhood-inferred label for a cell is different than its previous label, and if the ratio of confidence scores is above a threshold, we switch the old labels to the newly inferred label. We perform this operation synchronously for all cells and iterate for a fixed number of iterations or until it converges. As pointed out in previous work^48^, these naive approaches have the drawback of being influenced by the most dominant label in the network. To account for that, we have built a random model that assesses the total number of observed labels for each class within the neighborhood of cells, compared to their null distribution across the whole graph, as well as edge weights connecting cells to their neighbors. We use these computed p-value bounds in our framework instead of raw/adjusted frequencies of labels.

### Automatic cell state annotation

Given a continuous measurement at the cell level, we infer its enrichment at the cell state level using a permutation test. More specifically, we assess the co-concentration of cell measurements with the cell state encoding. For qualitative measurements, such as cell types, we construct its one-hot-encoding and similarly perform a permutation test.

## Supplementary Figure Legends

**Supplementary Figure 1.**
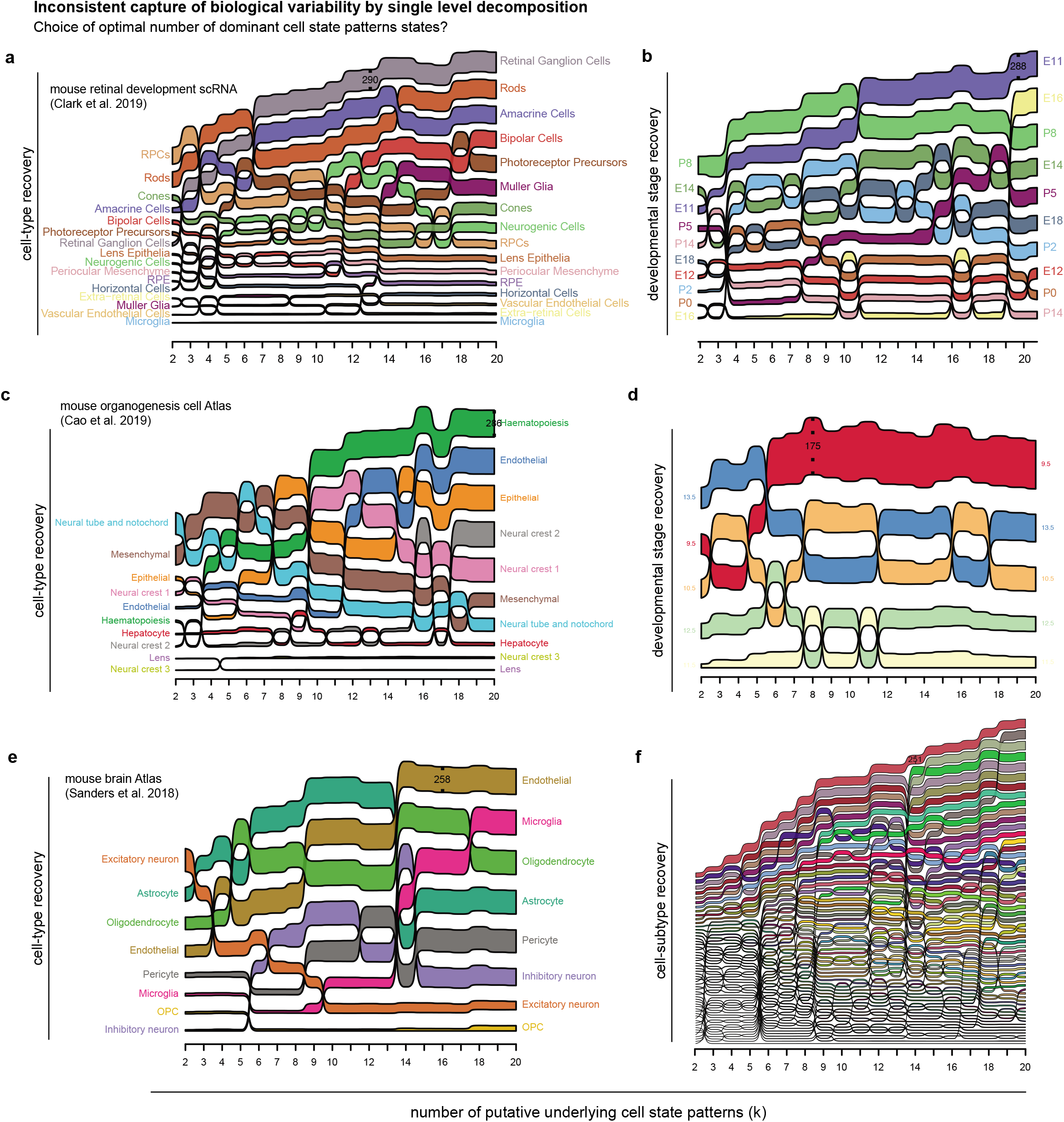
Dependency of cell type, lineage, and trajectory capture efficiency on the number of latent cell states. **(a-b)** whole tissue development (mouse retina development). Some cell types/stages, such as RGC/E11, show a significant enhancement in capture rate after increasing the total number of archetypes, while for others, such as RPCs, they show degradation in performance. **(c-d)** whole organ development (mouse organogenesis). Similar patterns emerge showing a heterogeneous pattern of variable capture rate changes as a function of latent cell states. **(e-f)** whole tissue atlas (mouse brain).

**Supplementary Figure 2.**
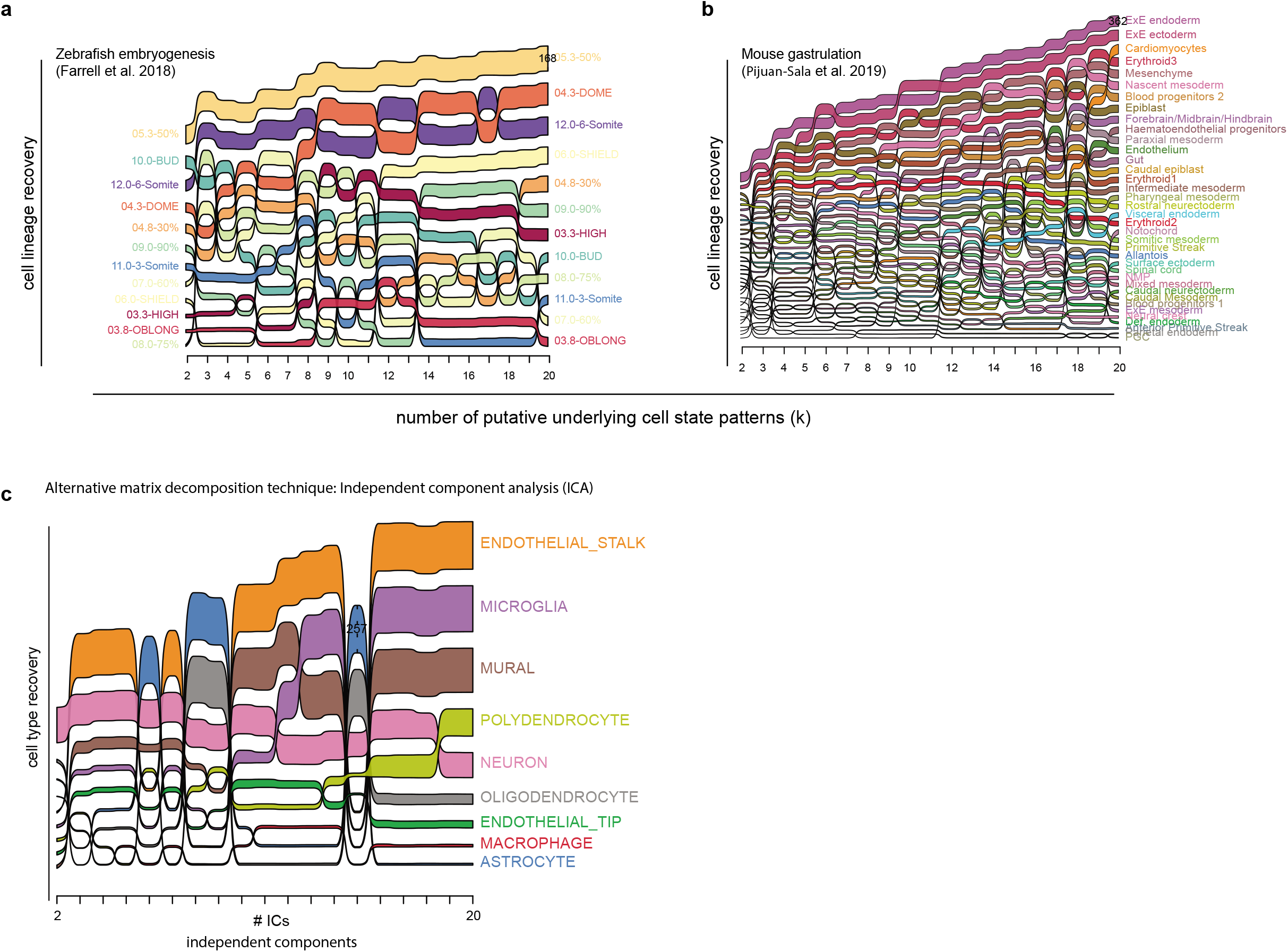
Additional support for heterogeneity of capture efficiency. **(a-b)** Additional time-series datasets, showing consistency of dependency patterns. **(c)** Similar pattern emerges when looking at alternative decomposition techniques (ICA).

**Supplementary Figure 3.**
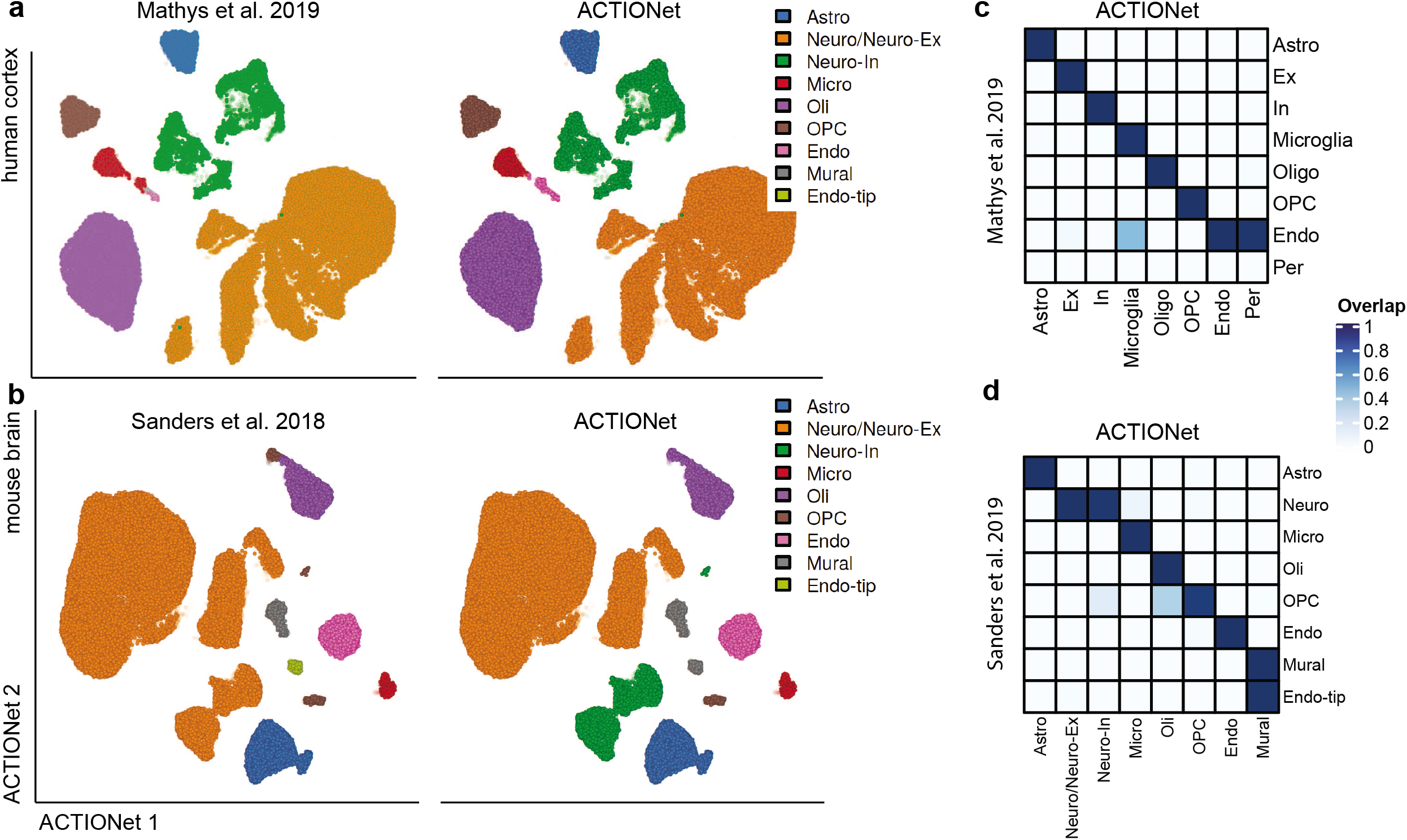
Performance of automated cell-type annotation. Reported (left) and inferred (right) cell type annotations for human (top) and mouse (bottom) brain PFC cells. **(a-c)** annotation consistency for cell types in the human brain PFC tissue. **(d-e)** annotation consistency for mouse PFC cortex. Annotations show overall consistency. Additionally, major neuron type distinction (inhibitory/excitatory), not reported in the original annotation, are automatically detected based on known marker genes.

**Supplementary Figure 4.**
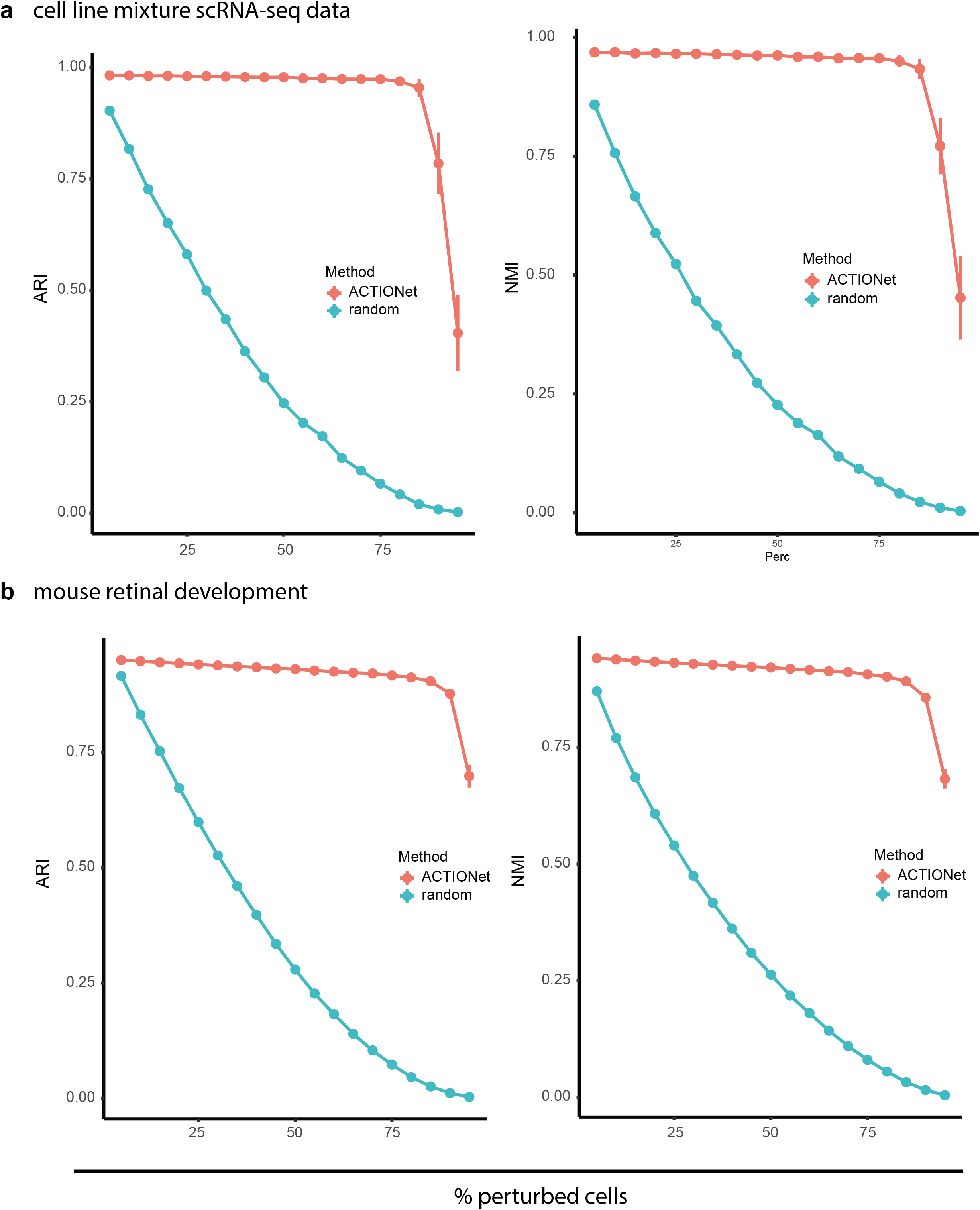
Performance of mislabel corrections. Adjusted rand index (ARI) (left) and Normalized Mutual Index (NMI) (right) cell types for synthetic cell line mixtures (top) and mouse retina development (bottom). X-axis is the percent of cells that their label has been perturbed, while Y-axis is the performance degradation upon perturbation (blue)/recovery upon correction (red).

